# Rare copy number variants (CNVs) and breast cancer risk

**DOI:** 10.1101/2021.05.20.444828

**Authors:** Joe Dennis, Jonathan P. Tyrer, Logan C. Walker, Kyriaki Michailidou, Leila Dorling, Manjeet K. Bolla, Qin Wang, Thomas U. Ahearn, Irene L. Andrulis, Hoda Anton-Culver, Natalia N. Antonenkova, Volker Arndt, Kristan J. Aronson, Laura E. Beane Freeman, Matthias W. Beckmann, Sabine Behrens, Javier Benitez, Marina Bermisheva, Natalia V. Bogdanova, Stig E. Bojesen, Hermann Brenner, Jose E. Castelao, Jenny Chang-Claude, Georgia Chenevix-Trench, Christine L. Clarke, NBCS Collaborators, J. Margriet Collée, CTS Consortium, Fergus J. Couch, Angela Cox, Simon S. Cross, Kamila Czene, Peter Devilee, Thilo Dörk, Laure Dossus, A. Heather Eliassen, Mikael Eriksson, D. Gareth Evans, Peter A. Fasching, Jonine Figueroa, Olivia Fletcher, Henrik Flyger, Lin Fritschi, Marike Gabrielson, Manuela Gago-Dominguez, Montserrat García-Closas, Graham G. Giles, Anna González-Neira, Pascal Guénel, Eric Hahnen, Christopher A. Haiman, Per Hall, Antoinette Hollestelle, Reiner Hoppe, John L. Hopper, Anthony Howell, ABCTB Investigators, kConFab Investigators, Agnes Jager, Anna Jakubowska, Esther M. John, Nichola Johnson, Michael E. Jones, Audrey Jung, Rudolf Kaaks, Renske Keeman, Elza Khusnutdinova, Cari M. Kitahara, Yon-Dschun Ko, Veli-Matti Kosma, Stella Koutros, Peter Kraft, Vessela N. Kristensen, Katerina Kubelka-Sabit, Allison W. Kurian, James V. Lacey, Diether Lambrechts, Nicole L. Larson, Martha Linet, Alicja Lukomska, Arto Mannermaa, Siranoush Manoukian, Sara Margolin, Dimitrios Mavroudis, Roger L. Milne, Taru A. Muranen, Rachel A. Murphy, Heli Nevanlinna, Janet E. Olson, Håkan Olsson, Tjoung-Won Park-Simon, Charles M. Perou, Paolo Peterlongo, Dijana Plaseska-Karanfilska, Katri Pylkäs, Gad Rennert, Emmanouil Saloustros, Dale P. Sandler, Elinor J. Sawyer, Marjanka K. Schmidt, Rita K. Schmutzler, Rana Shibli, Ann Smeets, Penny Soucy, Melissa C. Southey, Anthony J. Swerdlow, Rulla M. Tamimi, Jack A. Taylor, Lauren R. Teras, Mary Beth Terry, Ian Tomlinson, Melissa A. Troester, Thérèse Truong, Celine M. Vachon, Camilla Wendt, Robert Winqvist, Alicja Wolk, Xiaohong R. Yang, Wei Zheng, Argyrios Ziogas, Jacques Simard, Alison M. Dunning, Paul D.P. Pharoah, Douglas F. Easton

**Affiliations:** Centre for Cancer Genetic Epidemiology, Department of Public Health and Primary Care, University of Cambridge, Cambridge, UK; Centre for Cancer Genetic Epidemiology, Department of Oncology, University of Cambridge, Cambridge, UK; Department of Pathology and Biomedical Science, University of Otago, Christchurch, New Zealand; Biostatistics Unit, The Cyprus Institute of Neurology & Genetics, Nicosia, Cyprus; Cyprus School of Molecular Medicine, The Cyprus Institute of Neurology & Genetics, Nicosia, Cyprus; Division of Cancer Epidemiology and Genetics, National Cancer Institute, National Institutes of Health, Department of Health and Human Services, Bethesda, MD, USA; Fred A. Litwin Center for Cancer Genetics, Lunenfeld-Tanenbaum Research Institute of Mount Sinai Hospital, Toronto, ON, Canada; Department of Molecular Genetics, University of Toronto, Toronto, ON, Canada; Department of Medicine, Genetic Epidemiology Research Institute, University of California Irvine, Irvine, CA, USA; N.N. Alexandrov Research Institute of Oncology and Medical Radiology, Minsk, Belarus; Division of Clinical Epidemiology and Aging Research, German Cancer Research Center (DKFZ), Heidelberg, Germany; Department of Public Health Sciences, and Cancer Research Institute, Queen’s University, Kingston, ON, Canada; Department of Gynecology and Obstetrics, Comprehensive Cancer Center Erlangen-EMN, University Hospital Erlangen, Friedrich-Alexander University Erlangen-Nuremberg (FAU), Erlangen, Germany; Division of Cancer Epidemiology, German Cancer Research Center (DKFZ), Heidelberg, Germany; Biomedical Network on Rare Diseases (CIBERER), Madrid, Spain; Human Cancer Genetics Programme, Spanish National Cancer Research Centre (CNIO), Madrid, Spain; Institute of Biochemistry and Genetics, Ufa Federal Research Centre of the Russian Academy of Sciences, Ufa, Russia; Department of Radiation Oncology, Hannover Medical School, Hannover, Germany; Gynaecology Research Unit, Hannover Medical School, Hannover, Germany; Copenhagen General Population Study, Herlev and Gentofte Hospital, Copenhagen University Hospital, Herlev, Denmark; Department of Clinical Biochemistry, Herlev and Gentofte Hospital, Copenhagen University Hospital, Herlev, Denmark; Faculty of Health and Medical Sciences, University of Copenhagen, Copenhagen, Denmark; Division of Preventive Oncology, German Cancer Research Center (DKFZ) and National Center for Tumor Diseases (NCT), Heidelberg, Germany; German Cancer Consortium (DKTK), German Cancer Research Center (DKFZ), Heidelberg, Germany; Oncology and Genetics Unit, Instituto de Investigacion Sanitaria Galicia Sur (IISGS), Xerencia de Xestion Integrada de Vigo-SERGAS, Vigo, Spain; Cancer Epidemiology Group, University Cancer Center Hamburg (UCCH), University Medical Center Hamburg-Eppendorf, Hamburg, Germany; Department of Genetics and Computational Biology, QIMR Berghofer Medical Research Institute, Brisbane, Queensland, Australia; Westmead Institute for Medical Research, University of Sydney, Sydney, New South Wales, Australia; Department of Cancer Genetics, Institute for Cancer Research, Oslo University Hospital-Radiumhospitalet, Oslo, Norway; Institute of Clinical Medicine, Faculty of Medicine, University of Oslo, Oslo, Norway; Department of Research, Vestre Viken Hospital, Drammen, Norway; Section for Breast- and Endocrine Surgery, Department of Cancer, Division of Surgery, Cancer and Transplantation Medicine, Oslo University Hospital-Ullevål, Oslo, Norway; Department of Radiology and Nuclear Medicine, Oslo University Hospital, Oslo, Norway; Department of Pathology, Akershus University Hospital, Lørenskog, Norway; Department of Tumor Biology, Institute for Cancer Research, Oslo University Hospital, Oslo, Norway; Department of Oncology, Division of Surgery, Cancer and Transplantation Medicine, Oslo University Hospital-Radiumhospitalet, Oslo, Norway; National Advisory Unit on Late Effects after Cancer Treatment, Oslo University Hospital, Oslo, Norway; Department of Oncology, Akershus University Hospital, Lørenskog, Norway; Oslo Breast Cancer Research Consortium, Oslo University Hospital, Oslo, Norway; Department of Medical Genetics, Oslo University Hospital and University of Oslo, Oslo, Norway; Department of Clinical Genetics, Erasmus University Medical Center, Rotterdam, The Netherlands; Department of Computational and Quantitative Medicine, City of Hope, Duarte, CA, USA; City of Hope Comprehensive Cancer Center, City of Hope, Duarte, CA, USA; Department of Laboratory Medicine and Pathology, Mayo Clinic, Rochester, MN, USA; Sheffield Institute for Nucleic Acids (SInFoNiA), Department of Oncology and Metabolism, University of Sheffield, Sheffield, UK; Academic Unit of Pathology, Department of Neuroscience, University of Sheffield, Sheffield, UK; Department of Medical Epidemiology and Biostatistics, Karolinska Institutet, Stockholm, Sweden; Department of Pathology, Leiden University Medical Center, Leiden, The Netherlands; Department of Human Genetics, Leiden University Medical Center, Leiden, The Netherlands; Nutrition and Metabolism Section, International Agency for Research on Cancer (IARC-WHO), Lyon, France; Channing Division of Network Medicine, Department of Medicine, Brigham and Women’s Hospital and Harvard Medical School, Boston, MA, USA; Department of Epidemiology, Harvard T.H. Chan School of Public Health, Boston, MA, USA; Division of Evolution and Genomic Sciences, School of Biological Sciences, Faculty of Biology, Medicine and Health, University of Manchester, Manchester Academic Health Science Centre, Manchester, UK; North West Genomics Laboratory Hub, Manchester Centre for Genomic Medicine, St Mary’s Hospital, Manchester University NHS Foundation Trust, Manchester Academic Health Science Centre, Manchester, UK; David Geffen School of Medicine, Department of Medicine Division of Hematology and Oncology, University of California at Los Angeles, Los Angeles, CA, USA; Usher Institute of Population Health Sciences and Informatics, The University of Edinburgh, Edinburgh, UK; Cancer Research UK Edinburgh Centre, The University of Edinburgh, Edinburgh, UK; The Breast Cancer Now Toby Robins Research Centre, The Institute of Cancer Research, London, UK; Department of Breast Surgery, Herlev and Gentofte Hospital, Copenhagen University Hospital, Herlev, Denmark; School of Public Health, Curtin University, Perth, Western Australia, Australia; Fundación Pública Galega de Medicina Xenómica, Instituto de Investigación Sanitaria de Santiago de Compostela (IDIS), Complejo Hospitalario Universitario de Santiago, SERGAS, Santiago de Compostela, Spain; Moores Cancer Center, University of California San Diego, La Jolla, CA, USA; Cancer Epidemiology Division, Cancer Council Victoria, Melbourne, Victoria, Australia; Centre for Epidemiology and Biostatistics, Melbourne School of Population and Global Health, The University of Melbourne, Melbourne, Victoria, Australia; Precision Medicine, School of Clinical Sciences at Monash Health, Monash University, Clayton, Victoria, Australia; Center for Research in Epidemiology and Population Health (CESP), Team Exposome and Heredity, INSERM, University Paris-Saclay, Villejuif, France; Center for Familial Breast and Ovarian Cancer, Faculty of Medicine and University Hospital Cologne, University of Cologne, Cologne, Germany; Center for Integrated Oncology (CIO), Faculty of Medicine and University Hospital Cologne, University of Cologne, Cologne, Germany; Department of Preventive Medicine, Keck School of Medicine, University of Southern California, Los Angeles, CA, USA; Department of Oncology, Södersjukhuset, Stockholm, Sweden; Department of Medical Oncology, Erasmus MC Cancer Institute, Rotterdam, The Netherlands; Dr. Margarete Fischer-Bosch-Institute of Clinical Pharmacology, Stuttgart, Germany; University of Tübingen, Tübingen, Germany; Division of Cancer Sciences, University of Manchester, Manchester, UK; Australian Breast Cancer Tissue Bank, Westmead Institute for Medical Research, University of Sydney, Sydney, New South Wales, Australia; Research Department, Peter MacCallum Cancer Center, Melbourne, Victoria, Australia; Sir Peter MacCallum Department of Oncology, The University of Melbourne, Melbourne, Victoria, Australia; Department of Genetics and Pathology, Pomeranian Medical University, Szczecin, Poland; Independent Laboratory of Molecular Biology and Genetic Diagnostics, Pomeranian Medical University, Szczecin, Poland; Department of Epidemiology & Population Health, Stanford University School of Medicine, Stanford, CA, USA; Department of Medicine, Division of Oncology, Stanford Cancer Institute, Stanford University School of Medicine, Stanford, CA, USA; Division of Genetics and Epidemiology, The Institute of Cancer Research, London, UK; Division of Molecular Pathology, The Netherlands Cancer Institute - Antoni van Leeuwenhoek Hospital, Amsterdam, The Netherlands; Saint Petersburg State University, Saint Petersburg, Russia; Radiation Epidemiology Branch, Division of Cancer Epidemiology and Genetics, National Cancer Institute, Bethesda, MD, USA; Department of Internal Medicine, Evangelische Kliniken Bonn gGmbH, Johanniter Krankenhaus, Bonn, Germany; Translational Cancer Research Area, University of Eastern Finland, Kuopio, Finland; Institute of Clinical Medicine, Pathology and Forensic Medicine, University of Eastern Finland, Kuopio, Finland; Biobank of Eastern Finland, Kuopio University Hospital, Kuopio, Finland; Program in Genetic Epidemiology and Statistical Genetics, Harvard T.H. Chan School of Public Health, Boston, MA, USA; Department of Histopathology and Cytology, Clinical Hospital Acibadem Sistina, Skopje, Republic of North Macedonia; VIB Center for Cancer Biology, Leuven, Belgium; Laboratory for Translational Genetics, Department of Human Genetics, University of Leuven, Leuven, Belgium; Department of Health Sciences Research, Mayo Clinic, Rochester, MN, USA; Unit of Medical Genetics, Department of Medical Oncology and Hematology, Fondazione IRCCS Istituto Nazionale dei Tumori di Milano, Milan, Italy; Department of Clinical Science and Education, Södersjukhuset, Karolinska Institutet, Stockholm, Sweden; Department of Medical Oncology, University Hospital of Heraklion, Heraklion, Greece; Department of Obstetrics and Gynecology, Helsinki University Hospital, University of Helsinki, Helsinki, Finland; School of Population and Public Health, University of British Columbia, Vancouver, BC, Canada; Cancer Control Research, BC Cancer, Vancouver, BC, Canada; Department of Cancer Epidemiology, Clinical Sciences, Lund University, Lund, Sweden; Department of Genetics, Lineberger Comprehensive Cancer Center, University of North Carolina at Chapel Hill, Chapel Hill, NC, USA; Genome Diagnostics Program, IFOM - the FIRC Institute of Molecular Oncology, Milan, Italy; Research Centre for Genetic Engineering and Biotechnology ‘Georgi D. Efremov’, MASA, Skopje, Republic of North Macedonia; Laboratory of Cancer Genetics and Tumor Biology, Cancer and Translational Medicine Research Unit, Biocenter Oulu, University of Oulu, Oulu, Finland; Laboratory of Cancer Genetics and Tumor Biology, Northern Finland Laboratory Centre Oulu, Oulu, Finland; Clalit National Cancer Control Center, Carmel Medical Center and Technion Faculty of Medicine, Haifa, Israel; Department of Oncology, University Hospital of Larissa, Larissa, Greece; Epidemiology Branch, National Institute of Environmental Health Sciences, NIH, Research Triangle Park, NC, USA; School of Cancer & Pharmaceutical Sciences, Comprehensive Cancer Centre, Guy’s Campus, King’s College London, London, UK; Division of Psychosocial Research and Epidemiology, The Netherlands Cancer Institute - Antoni van Leeuwenhoek hospital, Amsterdam, The Netherlands; Center for Molecular Medicine Cologne (CMMC), Faculty of Medicine and University Hospital Cologne, University of Cologne, Cologne, Germany; Department of Surgical Oncology, University Hospitals Leuven, Leuven, Belgium; Genomics Center, Centre Hospitalier Universitaire de Québec – Université Laval Research Center, Québec City, QC, Canada; Department of Clinical Pathology, The University of Melbourne, Melbourne, Victoria, Australia; Division of Breast Cancer Research, The Institute of Cancer Research, London, UK; Department of Population Health Sciences, Weill Cornell Medicine, New York, NY, USA; Epigenetic and Stem Cell Biology Laboratory, National Institute of Environmental Health Sciences, NIH, Research Triangle Park, NC, USA; Department of Population Science, American Cancer Society, Atlanta, GA, USA; Department of Epidemiology, Mailman School of Public Health, Columbia University, New York, NY, USA; Institute of Cancer and Genomic Sciences, University of Birmingham, Birmingham, UK; Wellcome Trust Centre for Human Genetics and Oxford NIHR Biomedical Research Centre, University of Oxford, Oxford, UK; Department of Epidemiology, Gillings School of Global Public Health and UNC Lineberger Comprehensive Cancer Center, University of North Carolina at Chapel Hill, Chapel Hill, NC, USA; Department of Quantitative Health Sciences, Division of Epidemiology, Mayo Clinic, Rochester, MN, USA; Institute of Environmental Medicine, Karolinska Institutet, Stockholm, Sweden; Department of Surgical Sciences, Uppsala University, Uppsala, Sweden; Division of Epidemiology, Department of Medicine, Vanderbilt Epidemiology Center, Vanderbilt-Ingram Cancer Center, Vanderbilt University School of Medicine, Nashville, TN, USA

## Abstract

**Background:** Copy number variants (CNVs) are pervasive in the human genome but potential disease associations with rare CNVs have not been comprehensively assessed in large datasets. We analysed rare CNVs in genes and non-coding regions for 86,788 breast cancer cases and 76,122 controls of European ancestry with genome-wide array data.

**Results:** Gene burden tests detected the strongest association for deletions in *BRCA1* (P= 3.7E-18). Nine other genes were associated with a p-value < 0.01 including known susceptibility genes *CHEK2* (P= 0.0008), *ATM* (P= 0.002) and *BRCA2* (P= 0.008). Outside the known genes we detected associations with p-values < 0.001 for either overall or subtype-specific breast cancer at nine deletion regions and four duplication regions. Three of the deletion regions were in established common susceptibility loci.

**Conclusions:** This is the first genome-wide analysis of rare CNVs in a large breast cancer case-control dataset. We detected associations with exonic deletions in established breast cancer susceptibility genes. We also detected suggestive associations with non-coding CNVs in known and novel loci with large effects sizes. Larger sample sizes will be required to reach robust levels of statistical significance.

## 1 Introduction

Copy number variants (CNVs) are pervasive in the human genome but are more challenging to detect with current technologies than single nucleotide variants (SNVs). Recent comprehensive sequencing projects ^1,2^ have characterised CNVs in large sample sets. The gnomAD project identified a median of 3,505 deletions and 723 duplications covering more than 50 base pairs per genome. Most deletions and duplications tend to be rare with longer variants tending to be rarer, suggesting negative selection against these variants. At the population level the 1000 Genomes project has mapped a large proportion of inherited CNVs ^3^ and observed that 65% had a frequency below 2%.

Some CNVs have an established role in the inherited risk of breast cancer. Rare loss of function variants in susceptibility genes such as *BRCA1* and *CHEK2* are associated with a large increase in risk^4^. While the majority of these variants are single nucleotide mutations and short indels, they also include longer deletions and duplications. It has been reported that up to a third of loss of function *BRCA1* variants in some populations may be CNVs ^5^.

Large-scale genome-wide association studies (GWAS) have established breast cancer associations with common variants at more than 150 loci, mostly in non-coding regions^6–9^. At two of the loci, deletions imputed from the 1000 Genomes reference panel have been identified as likely causal variants. A deletion of the *APOBEC3B* gene-coding region increases breast cancer risk^10^ and analysis of the tumours of the germline deletion carriers showed an increase in APOBEC-mediated somatic mutations.^11^ A deletion in a regulatory region was identified as a likely causal variant at the 2q35 locus^12,13^.

Detecting CNVs from the intensity measurements of genotyping array probes is prone to producing unreliable calls due to the high level of noise. We recently developed a novel CNV calling method, CamCNV^14^, which focuses on rare CNVs and identifies outlier samples that may have a CNV, based on the intensity distribution across all samples at each probe. We showed that this approach is able to detect CNVs using as few as three probes^14^. Here, we apply this approach to a very large array genotype dataset to search for novel breast cancer associated CNVs.

## 2 Data

### 2.1 Subjects

Data were derived from study participants in 66 studies participating in the Breast Cancer Association Consortium (BCAC) and genotyped as part of the OncoArray^7,15^ and iCOGS^6^ collaborations (Supplementary Table 1). Studies included population-based and hospitalbased case-control studies, and case-control studies nested within prospective cohorts; we only included data from studies that provided both cases and controls. Phenotype data were based on version 12 of the BCAC database. Cases were diagnosed with either invasive breast cancer or carcinoma-in-situ. Oestrogen receptor (ER) status was determined from medical records or tissue microarray evaluation, where available. Analyses were restricted to participants of European ancestry, as defined by ancestry informative principal components^6,7^. Where samples were genotyped on both arrays, we excluded the iCOGS sample as the OncoArray has better genome-wide coverage. After sample quality control (see below), data on 36,980 cases and 34,706 controls with iCOGS genotyping, and 49,808 cases and 41,416 controls with OncoArray genotyping, were available for analysis (Supplementary Table 2).

### 2.2 Arrays

The Illumina iCOGs genotyping array^6^ includes 211,155 probes for SNVs and short insertions/deletions. Most variants were selected because of previous association in casecontrol studies for breast prostate and ovarian cancers, or for dense mapping of regions harbouring an association. The OncoArray includes 533,631 probes, of which approximately half were selected from the Illumina HumanCore backbone, a set of SNPs designed to tag most common variants. The remainder were selected on the basis of evidence of previous association with breast, prostate, ovarian, lung or colorectal cancer risk. Approximately 32,000 variants on the OncoArray were selected to provide dense coverage of associated loci and known genes. The remainder were mostly selected from lists of common variants ranked by p-value, with a small number from lists of candidate variants.

## 3 Methods

### 3.1 CNV Calling

CNVs were called using the CamCNV pipeline as previously described^14^. In brief, the log R (LRR) intensity measurements and B allele frequency (BAF) for each sample at each probe were exported from Illumina’s Genome Studio software. A principal component adjustment (PCA) was applied to the LRR, grouped by study, to remove noise and batch effects. After removing noisy probes and those in regions with known common CNVs, the LRRs for each probe were converted to z-scores using the mean and standard deviation from all BCAC samples. Circular binary segmentation was applied to the z-scores ordered by probe position for each sample using the DNACopy package. ^16^ This produces a list of segments for each sample by chromosome where the z-score of consecutive probes changes by more than two standard deviations. Segments with a mean probe z-score between −3.7 and −14 were called as deletions and segments with a mean z-score between +2 and +10 as duplications. We restricted the calls to segments covering a minimum of three and a maximum of 200 probes.

As per the CamCNV pipeline, we then excluded deletions with inconsistent B Allele Frequency and CNVs with a shift in LRR at the sample level that was outside the expected range. The additional CNV exclusions are summarised in Supplementary Table 3. To exclude regions with a high level of noise we also excluded CNVs falling within 1Mb of telomeres and centromeres and a number of immune loci such as the T-cell receptor genes where somatic mutations in blood are often observed ^17^.

### 3.2 Sample Quality Control

Standard sample quality control exclusions were performed, as previously described for the SNP genotype analyses^6,7^. These include exclusions for excess heterozygosity, ancestry outliers, mismatches with other genotyping, and close relatives. A stricter sample call rate of >99% was used for the CNV analysis, compared to >95% used in the genotype analyses. We also excluded any participants for whom a DNA sample was not collected from blood and any that had been whole genome amplified.

In addition, we used two metrics to exclude noisy samples liable to produce an excess of unreliable CNV calls. First, we calculated a derivative log ratio spread (DLRS) figure for each sample as the standard deviation of the differences between LRR for probes ordered by genomic position, divided by the square root of two. This measures the variance in the LRR from each probe to the next averaged over the whole genome and thus is insensitive to large fluctuations such as might be expected between different chromosomes in the same sample. An ideal sample would have a small DLRS as the only variance would come from a small number of genuine CNVs. We calculated the DLRS using the dLRs function in R package ADM3 (https://CRAN.R-project.org/package=ADM3) before and after the PCA. At both stages we excluded samples with a DLRS more than 3.5 standard deviations above the mean DLRS for that study.

Second, we counted the number of short segments (between three and 200 probes) output by DNACopy for each sample. We observed that the distribution of segment counts was skewed to the right with an excess of samples with a large number of segments. We calculated a cut-off for the maximum number of segments using the following formula where x is the segment count for each sample (based on the rationale that the distribution of the true number of segments should be approximately Poisson):

y=2*sqrt(x)
cut-off = median(y)+3.5

The sample exclusions resulting from these QC steps are summarised in Supplementary Table 4.

### 3.3 Association Tests

All analyses were carried out separately for deletions and duplications. As we were only assessing rare CNVs, we treated all carriers as heterozygotes and did not attempt to identify rare homozygotes.

To account for overlapping CNVs and imprecision in the breakpoints, we assigned individual CNVs to regions. To identify the regions, we moved sequentially along each chromosome, identifying the start as an Oncoarray probe position where deletions were observed in at least five samples, and then the end position as the probe position before the first probe where deletions were observed in fewer than five samples. Regions within five probes of each other were then merged together. The process was repeated for duplications. Regions were also merged such that the major susceptibility genes (*BRCA1*, *BRCA2*, *CHEK2*) were included within a single region. We then assigned individual CNVs to regions where at least 90% of the CNV’s length fell within the region. For iCOGS, which generally has less dense probe coverage, we first assigned CNVs to the OncoArray regions where they showed > 90% overlap. We then assigned any remaining CNVs to regions defined using the iCOGS probes, using the same procedure. Using this approach, 3,306 deletion regions were identified from OncoArray data, 812 of which were also observed using iCOGS data, and 541 regions identified using iCOGS alone. For duplications there were 2,203 OncoArray regions, with 854 also observed using iCOGS data, and 483 iCOGS specific regions.

Associations were evaluated for each array and each region using logistic regression, to derive a log odds ratio per deletion/duplication. Statistical significance was evaluated using a likelihood ratio test. The logistic regression analyses were conducted using in-house software (https://ccge.medschl.cam.ac.uk/software/mlogit/). Study and ten ancestry informative principal components, defined separately for each array, were included as covariates. The results from each array were combined in a standard fixed effect metaanalysis using the METAL software^18^. To avoid regions with too few observations, we excluded regions with fewer than 24 deletions or duplications (~0.015% of samples).

To detect more precisely the location of association signals, we also generated results for each probe. We created a vector of pseudo-genotypes for each probe with samples, such that a deletion covering that probe was coded as 1 and all other samples were coded as 0. We generated a similar set of genotypes for duplications. The results were analysed using logistic regression, as above.

To test for association between CNVs affecting the coding sequence of genes, in aggregate, and breast cancer risk, we identified samples with a deletion or duplication overlapping the exons of each gene. Exon positions were downloaded from the UCSC Genome Browser hg19 knownGene table. We used logistic regression to generate a log odds ratio (OR) for carriers of coding variants covering each gene, adjusted for study, as above. We generated results for each array and then for carriers combined across both arrays. For the combined analyses we treated studies with samples on both arrays as separate studies.

To calculate Bayesian False Discovery Probabilities (BFDPs) we assumed a log-normally distributed prior effect size as described by Wakefield^19^. The prior log(OR) was determined by assuming a 95% probability that the OR was less than some bound K, where K=3 for the regional and gene-based analysis, except for *BRCA1* and *BRCA2* where K=20 was assumed. The prior probability of association was assumed to be 0.001 for the regional analysis, 0.99 for *BRCA1*, *BRCA2*, *ATM* and *CHEK2* and 0.002 for other genes. For the gene-based analysis only positive associations were considered as the prior evidence for all genes was in favour of PTVs being positively associated with risk.

To determine whether there was a tendency for CNVs to be associated with an excess, or deficit, of risk across genes or regions, we computed signed z-scores as the square root of the chi-squared statistic for each gene, multiplied by ±1 depending on whether the effect estimate was positive or negative. These were ranked and normalised summed z-scores, based on the r most significant associations, were derived. The overall test statistic was the maximum summed z-score over all possible values of *r*:

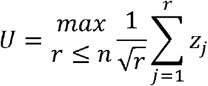

Where *n* is the total number of genes/regions being tested. The significance of U was then determined by permutation, randomly permuting case-control labels within study 50 times.

## 4 Results

### 4.1 Summary of CNVs Detected

After quality control we detected a mean of 2.9 deletions (standard deviation 1.6) and 2.5 duplications (SD 2.0) per sample. Supplementary Table 5 shows the mean length, probe coverage and segment z-scores of called CNVs. Duplications tended to be longer than deletions: for example, deletions called on OncoArray covered a mean of 45 Kilobases (Kb) (SD 106 Kb) over 9.8 probes (SD 17.2), while duplications covered a mean of 109 Kb (SD 202 Kb) over 18.9 probes (SD 36.5). CNV calls observed in multiple samples were concentrated in a small proportion of probes (Supplementary Table 6), with <11% of probes having frequency >0.01% and <2% of probes having frequency >0.5%.

We identified called CNVs which overlapped for at least 90% of their length with rare deletions and duplications (frequency <1%) identified by the 1000 Genomes Project (Supplementary Table 5). Forty-nine percent of OncoArray deletions and 47% of iCOGs deletions matched a 1000 Genomes Project variant while 29% of OncoArray duplications and 20% of iCOGs duplications matched. In total we identified CNVs closely matching 3,273 of the deletion variants published by the 1000 Genomes Project (~9% of total) and 1,255 of their duplication variants (~24% of the total).

### 4.2 CNVs Associated with Overall Risk

Association results were derived for 1,301 regions containing deletions and 992 regions containing duplications. QQ plots are shown in Figure 1A for deletions and 1B for duplications. There was no evidence for inflation in the test statistics for duplications (lambda=0.98; lambda_1000_=1.00) and minimal evidence for deletions (lambda = 1.11; lambda_1000_=1.00).

**Figure 1.**
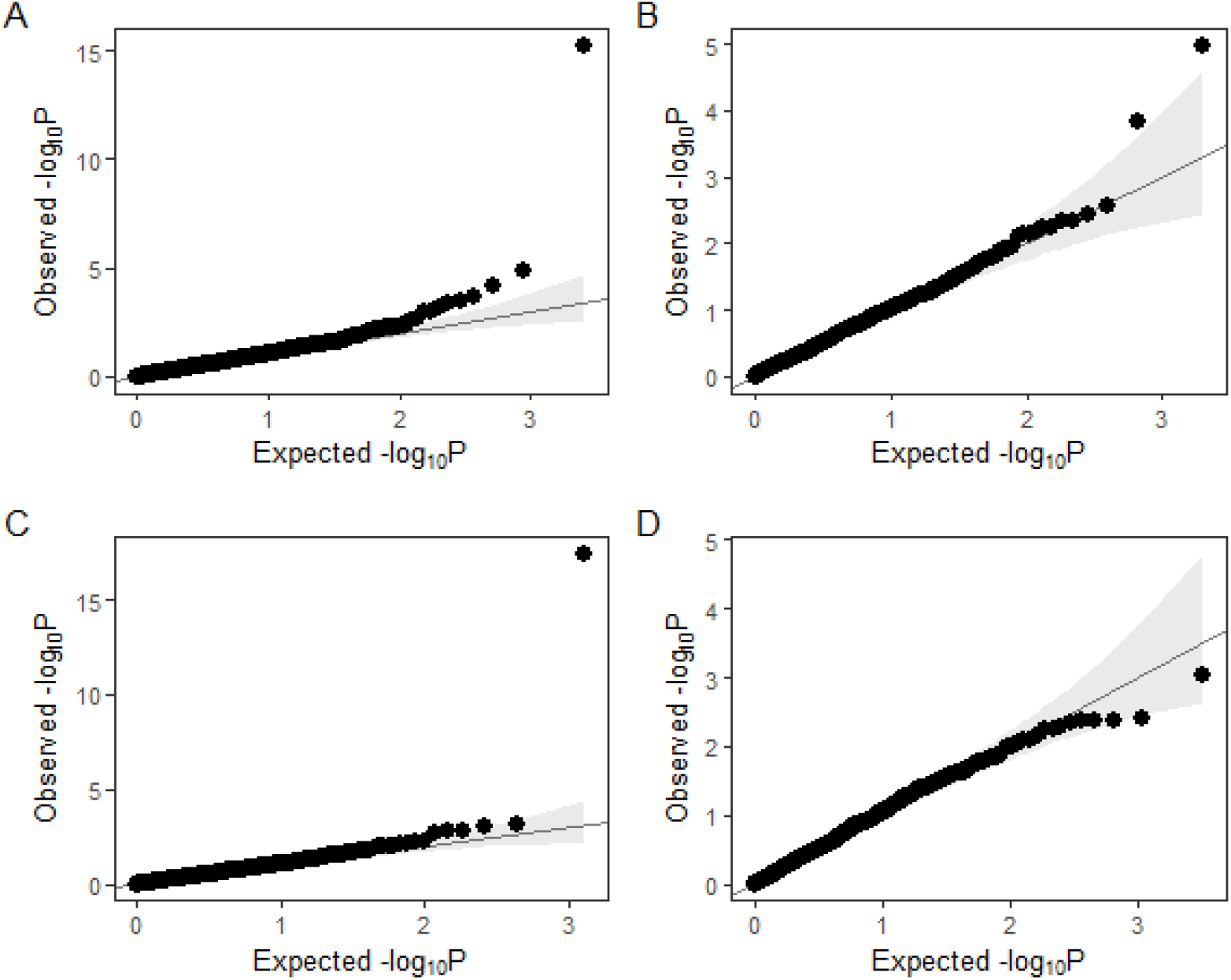
QQ plots for association of deletion regions (A), duplication regions (B), gene burden analysis for deletions (C) and gene burden for duplications (D).

Seven deletion and two duplication regions were associated with breast cancer risk at p<0.001 (Table 1); of these, deletions within the *BRCA1* region achieved p< 10^-6^. The results for all regions are shown in Supplementary Tables 7 and 8 and include statistics on the number of probes covered by the calls. The results for individual probes covered by the regions analysed are in Supplementary Tables 9 to 12.

**Table 1.** CNVs associated with overall risk

| Type | Locus | Chr. | Start<br>(Build37) | End | Total<br>Carriers | Odds<br>Ratio | Lower<br>CI | Upper<br>CI | Direction<br>(OncoArray,<br>iCOGs) | P-value |
| --- | --- | --- | --- | --- | --- | --- | --- | --- | --- | --- |
| Deletion | BRCA1 | 17 | 41188342 | 41363651 | 195 | 6.27 | 4.02 | 9.79 | ++ | 6.32E-16 |
| Deletion | Intergenic_FGFR2_ATE1 | 10 | 123435817 | 123461066 | 630 | 1.42 | 1.21 | 1.67 | ++ | 1.42E-05 |
| Deletion | Intergenic_ADCY8_EFR3A | 8 | 132199447 | 132250643 | 42 | 4.88 | 2.24 | 10.61 | ++ | 6.36E-05 |
| Deletion | KLHL1 | 13 | 70652321 | 71029916 | 1761 | 0.85 | 0.77 | 0.92 | -- | 2.31E-04 |
| Deletion | CHEK2 | 22 | 29083731 | 29123846 | 141 | 1.94 | 1.35 | 2.79 | ++ | 3.26E-04 |
| Deletion | SUPT3H | 6 | 44908728 | 45244478 | 32 | 0.23 | 0.1 | 0.52 | -? | 4.25E-04 |
| Deletion | Intergenic_GALNT1_C18orf21 | 18 | 33350917 | 33359197 | 123 | 1.92 | 1.32 | 2.78 | ?+ | 6.24E-04 |
| Duplication | 17p13.3_VPS53;NXN | 17 | 13905 | 1559829 | 577 | 0.69 | 0.59 | 0.82 | -- | 1.08E-05 |
| Duplication | 21q22.11_HUNK_LINC00159 | 21 | 33410933 | 33863246 | 102 | 2.23 | 1.47 | 3.38 | ++ | 1.48E-04 |

The *BRCA1* locus contains multiple deletions across the gene. The *CHEK2* region (OR 1.94, p=0.0003) covers the whole gene but nearly all the calls correspond to a deletion of exons nine and ten, which was previously observed in 1% of breast cancer cases and 0.4% of controls in Poland ^20^. We observed the deletion in 0.9% of Polish cases and 0.5% of controls.

The most significant association (OR=0.69 P=0.00001) for duplications covers a large region on 17p13.3 with multiple long variants overlapping shorter duplications. The OncoArray results by probe show the strongest associations at a series of probes (17: 814529-850542) in the first intron of *NXN*, with the lowest P-value at 17: 819141 (OR=0.45, P=0.002). The most significant probe position on iCOGs was also in this region (17:836631, OR=0.58, P=0.09) (Figure 2).

**Figure 2.**
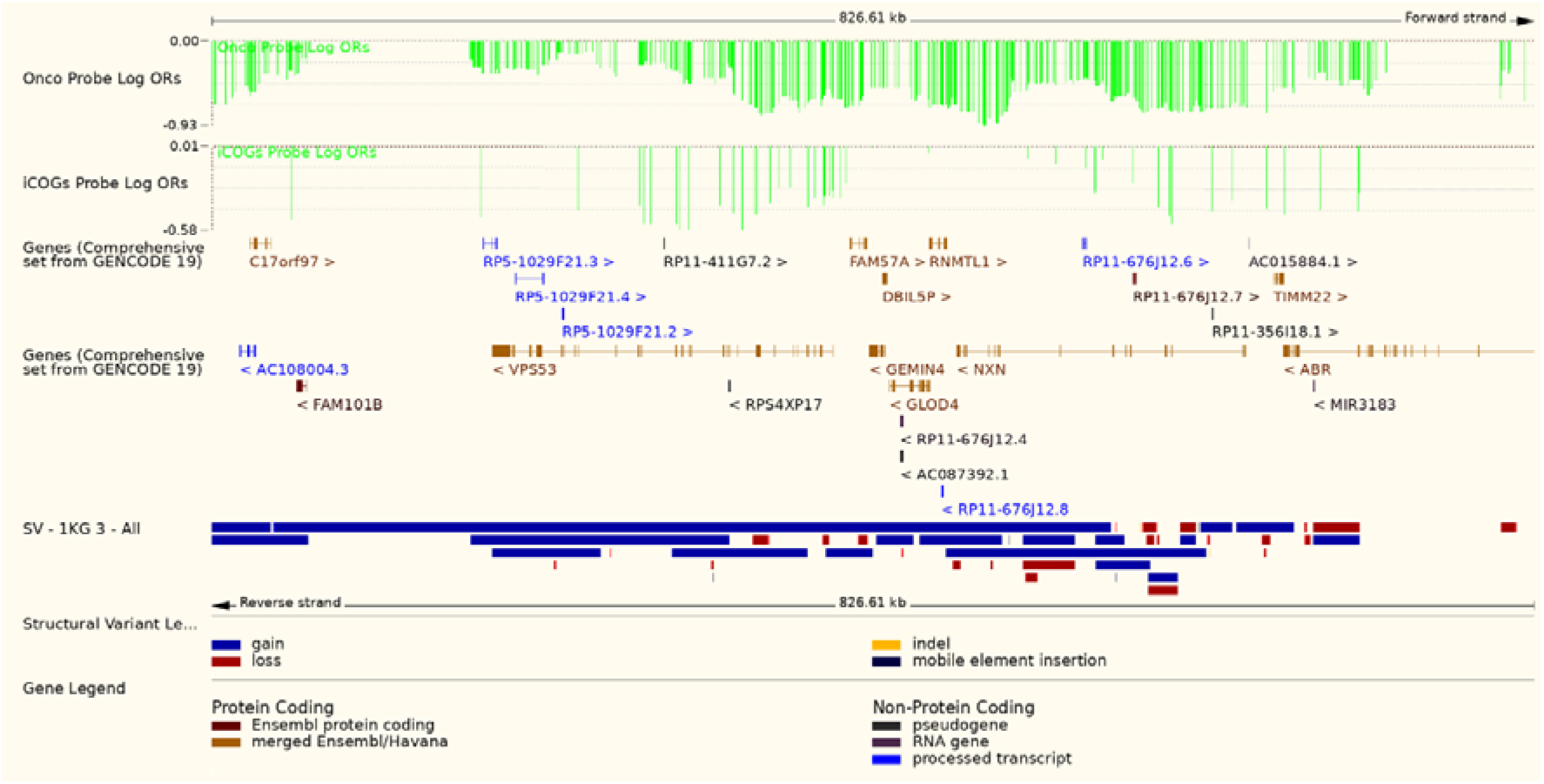
Plot of log odds ratios for probes within the 17p13.3 duplication region showing genes and 1000 Genomes CNVs from Ensembl.

### 4.3 Associations with Risk of Breast Cancer Subtypes

We repeated the analyses restricting cases to those with ER-positive, ER-negative and triple negative disease. Deletions and duplications with p-values below 0.001 are shown in Tables 2 and 3 and the results for all regions are in Supplementary Tables 13 and 14. An association was observed for *BRCA1* for all subtypes, with the exception of duplications for ER-positive disease. The odds ratio for *BRCA1* deletions was higher for ER-negative disease (OR=27.03; 95% CI, 15.66 to 46.67) than ER-positive (OR =2.81; CI, 1.56 to 5.08; P=8.46E-28 for the difference), while for *CHEK2* the odds ratio was higher for ER-positive disease (OR=2.32;CI,1.56 to 3.44) than ER-negative (OR=1.36; CI,0.66 to 2.82; P=0.11 for the difference), consistent with the known subtype-specific associations for deleterious variants in these genes ^21^.

**Table 2.** Subtype associations for deletions

| Subtype | Locus | Chr. | Start<br>(Build37) | End | Total<br>Carriers | Odds<br>Ratio | Lower<br>CI | Upper<br>CI | Direction<br>(OncoArray,<br>iCOGs) | P-value |
| --- | --- | --- | --- | --- | --- | --- | --- | --- | --- | --- |
| ER Positive | Intergenic_FGFR2_ATE1 | 10 | 123435817 | 123461066 | 478 | 1.52 | 1.27 | 1.81 | ++ | 5.04E-06 |
| ER Positive | CHEK2 | 22 | 29091788 | 29102967 | 79 | 2.36 | 1.56 | 3.44 | ++ | 3.03E-05 |
| ER Positive | ITGBL1 intronic | 13 | 102122905 | 102133221 | 58 | 3.29 | 1.83 | 5.92 | +? | 7.29E-05 |
| ER Positive | Intergenic_ADCY8_EFR3A | 8 | 132199447 | 132250643 | 31 | 4.95 | 2.20 | 11.30 | ++ | 1.07E-04 |
| ER Positive | BRCA1 | 17 | 41188342 | 41363651 | 57 | 2.81 | 1.55 | 5.08 | ++ | 6.41E-04 |
| ER Negative | BRCA1 | 17 | 41188342 | 41363651 | 112 | 27.03 | 15.66 | 46.67 | ++ | 2.62E-32 |
| ER Negative | Intergenic:ABCC4_CLDN10 | 13 | 95991263 | 96004144 | 134 | 2.16 | 1.43 | 3.26 | +? | 2.46E-04 |
| ER Negative | KLF12 intronic | 13 | 74356683 | 74357984 | 93 | 2.39 | 1.49 | 3.82 | +? | 2.89E-04 |
| ER Negative | Intergenic_DPP10_DDX18 | 2 | 118258797 | 118389164 | 44 | 3.34 | 1.64 | 6.80 | ++ | 8.84E-04 |
| Triple Neg. | BRCA1 | 17 | 41188342 | 41363651 | 72 | 40.55 | 21.70 | 75.76 | ++ | 3.64E-31 |
| Triple Neg. | Intergenic_DPP10_DDX18 | 2 | 118258797 | 118389164 | 40 | 6.56 | 2.83 | 15.18 | ++ | 1.13E-05 |
| Triple Neg. | Intergenic_DPP10_DDX18 | 2 | 117973154 | 118107795 | 48 | 4.54 | 1.88 | 10.97 | ++ | 7.92E-04 |

**Table 3.** Subtype associations for duplications

| Subtype | Locus | Chr. | Start<br>(Build37) | End | Total<br>Carriers | Odds<br>Ratio | Lower<br>CI | Upper<br>CI | Direction<br>(OncoArray,<br>iCOGs) | P-value |
| --- | --- | --- | --- | --- | --- | --- | --- | --- | --- | --- |
| ER Positive | 17p13.3_VPS53;NXN | 17 | 13905 | 1559829 | 454 | 0.67 | 0.55 | 0.81 | -- | 4.44E-05 |
| ER Positive | 21q22.11_HUNK_LINC00159 | 21 | 33410933 | 33863246 | 77 | 2.55 | 1.6 | 4.06 | ++ | 7.88E-05 |
| ER Positive | 15q13 | 15 | 28440287 | 32797352 | 1250 | 0.83 | 0.75 | 0.93 | -- | 6.95E-04 |
| ER Negative | BRCA1 | 17 | 41188342 | 41363651 | 42 | 5.93 | 2.31 | 15.19 | +- | 2.09E-04 |
| Triple Neg. | BRCA1 | 17 | 41188342 | 41363651 | 35 | 10.80 | 3.33 | 35.02 | +- | 7.29E-05 |
| Triple Neg. | Intergenic_TMC3_MEX3B | 15 | 81960409 | 82104822 | 231 | 2.39 | 1.55 | 3.71 | ++ | 9.25E-05 |

In total we observed five deletion and two duplication regions with p-values < 0.001 that were not below this threshold in the overall risk analysis. The strongest novel association for ER-positive was for an intronic deletion in *ITGBL1* (OR = 3.3, P=0.00007, P for difference by ER-status=0.18). For ER-negative disease the strongest novel association was with an intergenic deletion between *ABCC4 (MRP4)* and *CLDN10* (OR=2.16, P=0.0002, P for difference by ER-status=0.02). Neither of these associations was significant for the other subtype. For triple negative disease, the strongest evidence of association was found for an intergenic duplication between *TMC3* and *MEX3B* (OR=2.39 P= 0.00009) and for two separate deletions upstream of the *DDX18* gene: 2:118258797-118389164 (OR= 6.56, P= 0.00001) and 2:117973154-118107795 (OR=4.54, P= 0.0008). The association at these two deletions was driven by the same samples, with 75% of the carriers of the first deletion observed to have the second deletion and normal copy number at the 62kb gap between the deletions.

### 4.4 Associations at Established Common Susceptibility Loci

Three of the most significant associations were observed within regions harbouring known breast cancer susceptibility loci for breast cancer. The most significant (OR=1.42;CI,1.21 to 1.67; P=0.00015) was upstream of *FGFR2* and consistent with a 28 kb deletion in the 1000 Genomes Project data (chr10:123433204-123461492). Three independent risk signals have been previously identified at this region^22,23^. The effect size for the CNV was larger than those previously reported for these common SNPs (largest OR=1.27;CI,1.22 to 1.25). The CNV is in linkage disequilibrium with two of the SNPs identified as likely causally associated variants: rs35054928 (D’ = 0.82) and rs2981578 (D’ = 0.88). Conditioning on those SNPs reduced somewhat the strength of the association for the deletion (OR =1.30;CI 1.10 to 1.53;P=0.002, Supplementary Table 15).

The third strongest signal (OR=4.9 P=0.00001) in the deletion analysis for overall breast cancer was at 8: 132199447-132252439, 144Kb downstream of *ADCY8*. The strongest GWAS signal in this region lies in an intron of *ADCY8* (lead SNP rs73348588, OR =1.13, P= 8.2e-7)^7^. A 3kb deletion in intron 4 of *KLF12* was associated with ER-negative disease (OR = 2.4, P= 0.0007, P for difference by ER-status=0.01). This is 389kb distant from common SNPs, located between *KLF12* and *KLF5*, associated with ER-negative disease (rs9573140, OR = 0.94, P=3.62e-9) ^24^. The *KLF12* and *ADCY8* deletions are not in strong linkage disequilibrium with the corresponding GWAS signals and conditioning on these SNPs did not alter the strengths of the association for the CNVs (Supplementary Table 15).

### 4.5 Gene Burden tests

We performed gene burden tests based on CNVs that overlapped exons. Analyses were restricted to genes in which at least 24 CNVs were identified, leaving 645 genes with deletions (Supplementary Table 16) and 1596 genes with duplications (Supplementary Table 17). QQ plots are shown in Figures 1C for deletions and 1D for duplications. The lambda for inflation was 1.18 (; lambda_1000_=1.00) for deletions and 1.07 (; lambda_1000_=1.00) for duplications.

For deletions, we found 10 genes with P < 0.01 (Table 4), the most significant being *BRCA1* (OR=7.66, P= 3.72E-18). Four of these 10 genes (*ATM*, *BRCA1*, *BRCA2*, *CHEK2*) are known breast cancer susceptibility genes.^21^ Deletions were also observed in two other known susceptibility genes: *PALB2* (23 cases, 9 controls, OR=2.02, P=0.09) and *RAD51C* (21 cases, 9 controls, OR=2.04, P=0.08). The most significant novel association was for *SUPT3H* (OR=0.27, P=0.0004).

**Table 4.** Gene burden results for deletions

| Gene | Cases | Controls | Odds Ratio | Lower CI | Upper CI | P-value |
| --- | --- | --- | --- | --- | --- | --- |
| BRCA1 | 171 | 22 | 7.66 | 4.84 | 12.13 | 3.72E-18 |
| CHEK2 | 103 | 48 | 1.83 | 1.29 | 2.61 | 7.66E-04 |
| SUPT3H | 10 | 25 | 0.27 | 0.13 | 0.59 | 9.24E-04 |
| PCDHGB2 | 25 | 3 | 7.03 | 2.10 | 23.52 | 1.55E-03 |
| MEAK7 | 59 | 24 | 2.19 | 1.34 | 3.58 | 1.66E-03 |
| ATM | 35 | 8 | 3.43 | 1.56 | 7.51 | 2.11E-03 |
| MAD1L1 | 54 | 25 | 2.00 | 1.23 | 3.26 | 5.53E-03 |
| NPHP1 | 477 | 351 | 1.22 | 1.06 | 1.41 | 6.13E-03 |
| ZNF320 | 29 | 7 | 3.28 | 1.39 | 7.73 | 6.63E-03 |
| BRCA2 | 65 | 33 | 1.81 | 1.17 | 2.81 | 7.82E-03 |

For duplications we observed 15 genes with P < 0.01 (Table 5). The most significant association was for *VPS53* (OR = 0.5, P= 0.0009). This gene and *ABR* (OR=0.61 P= 0.008) both lie within the region on 17p which had the strongest association in the regional analysis. These associations were driven by duplications in different samples, with only one duplication in one sample overlapping both genes. Duplications were associated with an increased risk for only four of the 15 genes; the most statistically significant was *RSU1* (OR=3.4, P= 0.004). There was also some evidence of association with risk for duplications in *BRCA1* (OR = 1.75, P =0.01). However, analysis restricted to duplications that included exon 12 of *BRCA1* showed clearer evidence of association (34 carriers, OR = 4.7 P= 0.0001), consistent with one of the more frequent known *BRCA1* duplications that results in a frameshift^25^.

**Table 5.** Gene burden results for duplications

| Gene | Cases | Controls | Odds Ratio | Lower CI | Upper CI | P-value |
| --- | --- | --- | --- | --- | --- | --- |
| VPS53 | 40 | 65 | 0.50 | 0.33 | 0.75 | 9.46E-04 |
| ATP12A | 48 | 66 | 0.57 | 0.39 | 0.84 | 3.97E-03 |
| USP18 | 12 | 23 | 0.34 | 0.17 | 0.71 | 4.16E-03 |
| RPS6KA2 | 10 | 22 | 0.32 | 0.14 | 0.70 | 4.20E-03 |
| RSU1 | 21 | 8 | 3.40 | 1.47 | 7.84 | 4.23E-03 |
| AC008132.1 | 7 | 17 | 0.26 | 0.10 | 0.66 | 4.49E-03 |
| PNPLA4 | 479 | 346 | 1.23 | 1.06 | 1.42 | 5.30E-03 |
| NLGN4X | 291 | 320 | 0.79 | 0.67 | 0.93 | 5.55E-03 |
| ZNF439 | 31 | 9 | 2.98 | 1.37 | 6.45 | 5.72E-03 |
| TRIM6 | 5 | 19 | 0.24 | 0.09 | 0.67 | 6.34E-03 |
| RP11.363G10.2 | 13 | 27 | 0.39 | 0.20 | 0.78 | 7.39E-03 |
| USP31 | 7 | 18 | 0.30 | 0.12 | 0.73 | 8.02E-03 |
| TRDN | 15 | 29 | 0.42 | 0.22 | 0.80 | 8.26E-03 |
| ABR | 51 | 69 | 0.61 | 0.42 | 0.88 | 8.88E-03 |
| DNAJC15 | 109 | 67 | 1.52 | 1.11 | 2.09 | 9.29E-03 |

The gene burden subtype results are shown in Supplementary Tables 18 and 19. The strongest associations were observed for *BRCA1* deletions for ER Negative (OR = 33, P=5.5E-35) and Triple Negative (OR =49 P=7.1E-34) disease, *CHEK2* deletions for ER positive disease (OR = 2.14 P=0.0001) and *ATM* deletions for ER positive disease (OR=4.85, P=0.0001). No additional genes significant at P<0.0001 were found.

### 4.6 Direction of effect tests

In the gene burden and individual probe analyses we observed a directional effect, whereby the strongest associations for deletions tended to increase risk and those for duplications tended to be protective. To test whether these associations deviated from what would be expected by chance, we computed ranked summed z-score tests and evaluated the significance of the maximum test statistic by permutation. Results are summarised in Table 6. The statistic for deletion regions was more significant than any of the permuted statistics (P=0.04) but was reduced to P=0.12 after removing the known genes *BRCA1* and *CHEK2*. The significance of the gene burden test for deletions also was reduced from P=0.04 to P=0.2 when the known genes were removed. The statistic for the duplication regions was lower than any of the 50 permutations (P=0.04). The gene burden analysis for duplications was not significant.

**Table 6.** Direction of Effect Results

| Category | Analysis ustat | Max.ustat of 50 simulations | Min. ustat of 50 simulations | Simulations with larger/smaller ustat | P-value |
| --- | --- | --- | --- | --- | --- |
| Deletion regions | 9.48 | 6.96 | -7.81 | 0 | 0.04 |
| Deletion regions minus known<sup>1</sup> | 5.9 | 6.96 | -7.81 | 2 | 0.12 |
| Duplication regions | -9.2 | 6.54 | -6.99 | 0 | 0.04 |
| Gene Deletions | 9.18 | 5.64 | -9.67 | 0 | 0.04 |
| Gene Deletions minus known<sup>2</sup> | 5.01 | 5.64 | -9.67 | 4 | 0.20 |
| Gene Duplications | -4.33 | 5.96 | -11.26 | 33 | 1.29 |

1. BRCA1, CHEK2 regions excluded; 2. BRCA1, CHEK2, ATM, BRCA2 removed.

## 5 Discussion

We used the largest available breast cancer case-control dataset, comprising more than 86,000 cases and 76,000 controls with array genotyping, to test for associations with rare CNVs. Using the intensities from genotyping arrays to detect CNVs is not ideal due to a high level of noise and uncertainty in the calling, particularly for duplications. However, in tests of known CNVs and replication of calls between duplicate samples, the CamCNV method shows reasonable levels of sensitivity and specificity^14^. The main focus of this analysis was low frequency CNVs (<1% frequency) since higher frequency CNVs can generally be studied through imputation to a reference panel. In the 0.05%-1% frequency range, we could detect ~20% of the CNVs identified by the 1000 Genomes project. For some loci we only had evidence from one array because the probes do not exist to detect the variants on the other array. Thus, while this array-based approach provides power to evaluate the CNVs that can be assayed, much denser arrays or direct sequencing would be required to provide a complete evaluation of the contribution of CNVs.

In support of the reliability of the method, we detected evidence for both deletions and duplications in *BRCA1*, which was stronger for ER-negative disease, and for deletions in *CHEK2*, which were stronger for ER-positive disease. The latter appeared to be driven by a single founder deletion in East European populations. Weaker evidence of association was also observed for deletions in other susceptibility genes (*BRCA2*, *ATM*, *PALB2*, *RAD51C*); the ORs were consistent with those seen for deleterious SNVs and indels. ^21^ In total, around 0.5% of cases in our analysis had a deletion in one of the known susceptibility genes with the proportion rising to ~1% for cases diagnosed under 50 years of age. The majority of coding deletions are expected to affect only part of the gene, with one study observing that a quarter covered only a single exon.^26^ To detect all of these using array data would require at least three probes per exon. The OncoArray has this level of coverage for a few genes, including *BRCA1* and *BRCA2*, but the coverage is lower for most genes and many coding CNVs will have been missed.

A key issue is the appropriate level of statistical significance to apply to these analyses. For the gene burden tests, P<2.5×10^-6^, as used in exome-sequencing, seems an appropriate level. It is less clear what is appropriate for non-coding variants. A level of P<5×10^-8^ has become standard in GWAS and has been shown to lead to acceptable replication, but this seems over-conservative for CNVs, which are more likely to be deleterious. Consistent with this, for at least two of the ~200 common susceptibility loci, the likely causal variant is a CNV, a higher fraction than expected given the relative frequencies of CNVs and SNPs. Based on frequency analysis of whole-genome sequence data Abel et al. ^1^ estimated that rare CNVs are >800 times more likely to be deleterious than rare SNVs and >300 times more likely than rare indels. On the other hand, the significance level for non-coding CNVs should logically be more stringent than for the gene burden tests. Taken together, a significance level of ~10^-6^ seems appropriate, while associations at P<0.001 may be worth following up in future studies. In our analyses only the association at *BRCA1* (both in the overall and gene burden tests) passes the higher threshold. We also calculated Bayesian False Discovery Probabilities (BFDPs)^19^ (Supplementary Tables 20 and 21) for our associations using prior probabilities of 0.001 for regions and 0.002 for genes. Outside the known genes none of the BDFPs gave a probability below 10%, with the lowest BFDP of 0.11 for the deletion in the *FGFR2* locus. For a CNV observed with a frequency of 0.1% (n=91 samples in the OncoArray dataset) we had 40% power to detect an association with an odds ratio of 2 but only 1.5% power to detect an association with an odds ratio of 1.5. An OR of 2, comparable to that seen for deleterious variants in *ATM* and *CHEK2*, may be plausible for rare coding CNVs or non-coding CNVs that have a significant effect on gene expression. Larger sample sizes will clearly be required to evaluate rare CNVs with more modest associations.

In addition to the *BRCA1* and *CHEK2* loci, we found associations in three known susceptibility regions identified through GWAS, harbouring *FGFR2*, *ADCY8* and *KLF12*. In each case, the variants are rarer than the established associated variants, but confer higher risks. The *ADCY8* and *KLF12* deletions are not in linkage disequilibrium with the associated SNPs. The *FGFR2* deletion is in linkage disequilibrium with two of the likely causal common SNPs although there was still evidence of association with the deletion, albeit weaker, after conditional analysis. *In-silico* and functional analysis clearly demonstrate that FGFR2 is the target of the previously established variants^22,23^; it will be interesting to establish if the same is true for the CNV.

Excluding loci in known susceptibility regions, the strongest evidence of association was for a 12kb deletion (13:102121830-102133956) in the second intron of *ITGBL1* (OR = 3.3, P=0.00007 in the ER-positive analysis). This deletes a promoter flanking region (Ensembl ID: ENSR00001563823) and CTCF binding site (Ensembl ID: ENSR00001062398) active in mammary epithelial cells. There is experimental evidence that *ITGBL1* expression, mediated by the RUNX2 transcription factor, enables breast cancer cells to form bone metastases^27^.

In the gene burden analysis, the strongest novel association was for deletions within *SUPT3H*, which were associated with a reduced risk. *SUPT3H* encodes human SPT3, a component of the STAGA complex which acts as a co-activator of the MYC oncoprotein^28^. *SUPTH* is located within the first intron of the *RUNX2* transcription factor and the syntenic relationship between the two genes is highly evolutionarily conserved ^29^. *RUNX2* has a role in mammary gland development and high *RUNX2* expression is found in ER-negative tumours.^30^ The *PCDHGB2* association appears to be due to a single variant (5:140739812-140740918) that deletes the first exon but as this gene is part of the protocadherin gamma gene cluster it is also possible that the deletion may be having an effect on one of the five genes that overlap *PCDHGB2*. It also deletes a promoter active in mammary epithelial cells (ENSR00001342785). The next strongest signals were for *MEAK7* (OR=2.19 P= 0.001), a gene implicated in a mTOR signalling pathway^31^, and *MAD1L1* ((OR=2.00 P=0.005), a component of the mitotic spindle-assembly that has been suggested as a possible tumour suppressor^32^.

After *BRCA1*, the most significant association for ER Negative disease in the gene burden analysis was for *CYP2C18* which overlaps *CYP2C19* (ER-negative OR=2.6, P=0.002; triplenegative OR=4.4, P=0.0002). A previous analysis of CNVs and breast cancer in the Finnish population identified a founder mutation reaching an overall frequency of ~ 3% and reported a possible association at this locus for triple negative (OR 2.8, p=□0.02) and ER-negative breast cancer (OR =2.2 p=0.048).^33^

The results from duplications are harder to interpret as there are often longer duplications overlapping whole genes with shorter variants covering some part of their length. For the gene burden analysis there was little evidence of strong associations. In the regional analysis, the two strongest associations cover multiple genes. The strongest evidence of association (OR=0.69 P=1.1E-05) was for a 1.5 Mb region at the start of chromosome 17 (17:13905-1559829). The probe-specific and gene burden results highlighted some stronger signals within this region, for example within *NXN* and *VPS53*, but the direction of effect was consistent across the region with 80% of the OncoArray probes having an odds ratio of 0.75 or lower (Figure 2). For the 0.4Mb duplication region on chromosome 21 (OR= 2.23 P=0.0001) the probe-specific results from OncoArray highlighted that the signal is specific to a shorter intergenic region (21:33421860-33459975) between *HUNK* and *LINC00159*.

We observed some evidence of an aggregate directional effect, both for genes and non-genic regions, such that the deletions in aggregate were associated with increased risk. There was also some suggestion that duplications, in aggregate, were associated with a reduced risk. These results suggest that additional associations are present that could be established with a larger dataset. A new GWAS, Confluence (https://dceg.cancer.gov/research/cancer-types/breast-cancer/confluence-project), aims to double the available sample size for breast cancer. This GWAS includes probes specifically designed to assay some of the most significant CNVs observed in this study (those significant at P<0.001), and the sample size should be sufficient to confirm or refute these associations.

## Availability of Data and Materials

Code for the CamCNV calling algorithm and a test dataset with OncoArray genotyping is available at the project home page (https://github.com/jgd29/CamCNV). Full summary statistics for the regions and probes analysed are available in the Supplementary Tables. The majority of the OncoArray dataset analyzed in this study is available in the dbGap repository, Study ID: phs001265.v1.p1 (https://www.ncbi.nlm.nih.gov/projects/gap/cgi-bin/study.cgi?study_id=phs001265.v1.p1). The iCOGS dataset and complete OncoArray dataset cannot be made publicly available due to restraints imposed by the ethics committees of individual studies; requests for data can be made to the corresponding author or the Data Access Coordination Committee (DACC) of BCAC (http://bcac.ccge.medschl.cam.ac.uk/).

## Acknowledgements

We thank all the individuals who took part in these studies and all the researchers, clinicians, technicians and administrative staff who have enabled this work to be carried out. The COGS study would not have been possible without the contributions of the following Andrew Berchuck (OCAC), Rosalind A. Eeles, Ali Amin Al Olama, Zsofia Kote-Jarai, Sara Benlloch (PRACTICAL), Antonis Antoniou, Lesley McGuffog and Ken Offit (CIMBA), Andrew Lee, and Ed Dicks, Craig Luccarini and the staff of the Centre for Genetic Epidemiology Laboratory, the staff of the CNIO genotyping unit, Jacques Simard and Daniel C. Tessier, Francois Bacot, Daniel Vincent, Sylvie LaBoissière and Frederic Robidoux and the staff of the McGill University and Génome Québec Innovation Centre, Sune F. Nielsen, Borge G. Nordestgaard, and the staff of the Copenhagen DNA laboratory, and Julie M. Cunningham, Sharon A. Windebank, Christopher A. Hilker, Jeffrey Meyer and the staff of Mayo Clinic Genotyping Core Facility. ABCFS thank Maggie Angelakos, Judi Maskiell, Gillian Dite. ABCS thanks the Blood bank Sanquin, The Netherlands. ABCTB Investigators: Christine Clarke, Deborah Marsh, Rodney Scott, Robert Baxter, Desmond Yip, Jane Carpenter, Alison Davis, Nirmala Pathmanathan, Peter Simpson, J. Dinny Graham, Mythily Sachchithananthan. Samples are made available to researchers on a non-exclusive basis. BBCS thanks Eileen Williams, Elaine Ryder-Mills, Kara Sargus. BCEES thanks Allyson Thomson, Christobel Saunders, Terry Slevin, BreastScreen Western Australia, Elizabeth Wylie, Rachel Lloyd. The BCINIS study would not have been possible without the contributions of Dr. K. Landsman, Dr. N. Gronich, Dr. A. Flugelman, Dr. W. Saliba, Dr. F. Lejbkowicz, Dr. E. Liani, Dr. I. Cohen, Dr. S. Kalet, Dr. V. Friedman, Dr. O. Barnet of the NICCC in Haifa, and all the contributing family medicine, surgery, pathology and oncology teams in all medical institutes in Northern Israel. BIGGS thanks Niall McInerney, Gabrielle Colleran, Andrew Rowan, Angela Jones. The BREOGAN study would not have been possible without the contributions of the following: Angel Carracedo, Victor Muñoz Garzón, Alejandro Novo Domínguez, Maria Elena Martinez, Sara Miranda Ponte, Carmen Redondo Marey, Maite Peña Fernández, Manuel Enguix Castelo, Maria Torres, Manuel Calaza (BREOGAN), José Antúnez, Máximo Fraga and the staff of the Department of Pathology and Biobank of the University Hospital Complex of Santiago-CHUS, Instituto de Investigación Sanitaria de Santiago, IDIS, Xerencia de Xestion Integrada de Santiago-SERGAS; Joaquín González-Carreró and the staff of the Department of Pathology and Biobank of University Hospital Complex of Vigo, Instituto de Investigacion Biomedica Galicia Sur, SERGAS, Vigo, Spain. CBCS thanks study participants, co-investigators, collaborators and staff of the Canadian Breast Cancer Study, and project coordinators Agnes Lai and Celine Morissette. CCGP thanks Styliani Apostolaki, Anna Margiolaki, Georgios Nintos, Maria Perraki, Georgia Saloustrou, Georgia Sevastaki, Konstantinos Pompodakis. CGPS thanks staff and participants of the Copenhagen General Population Study. For the excellent technical assistance: Dorthe Uldall Andersen, Maria Birna Arnadottir, Anne Bank, Dorthe Kjeldgård Hansen. The Danish Cancer Biobank is acknowledged for providing infrastructure for the collection of blood samples for the cases. CNIO-BCS thanks Guillermo Pita, Charo Alonso, Nuria Álvarez, Pilar Zamora, Primitiva Menendez, the Human Genotyping-CEGEN Unit (CNIO). Investigators from the CPS-II cohort thank the participants and Study Management Group for their invaluable contributions to this research. They also acknowledge the contribution to this study from central cancer registries supported through the Centers for Disease Control and Prevention National Program of Cancer Registries, as well as cancer registries supported by the National Cancer Institute Surveillance Epidemiology and End Results program. The authors would like to thank the California Teachers Study Steering Committee that is responsible for the formation and maintenance of the Study within which this research was conducted. A full list of California Teachers Study team members is available at https://www.calteachersstudy.org/team. ESTHER thanks Hartwig Ziegler, Sonja Wolf, Volker Hermann, Christa Stegmaier, Katja Butterbach. FHRISK thanks NIHR for funding and the Manchester NIHR Biomedical Research Centre (IS-BRC-1215-20007). GC-HBOC thanks Stefanie Engert, Heide Hellebrand, Sandra Kröber and LIFE - Leipzig Research Centre for Civilization Diseases (Markus Loeffler, Joachim Thiery, Matthias Nüchter, Ronny Baber). The GENICA Network: Dr. Margarete Fischer-Bosch-Institute of Clinical Pharmacology, Stuttgart, and University of Tübingen, Germany [Hiltrud Brauch, RH, Wing-Yee Lo], German Cancer Consortium (DKTK) and German Cancer Reseach Center (DKFZ), Partner Site Tübingen, 72074 Tübingen, Germany [Hiltrud Brauch], gefördert durch die Deutsche Forschungsgemeinschaft (DFG) im Rahmen der Exzellenzstrategie des Bundes und der Länder – EXC 2180 – 390900677 [Hiltrud Brauch], Department of Internal Medicine, Evangelische Kliniken Bonn gGmbH, Johanniter Krankenhaus, Bonn, Germany [YDK, Christian Baisch], Institute of Pathology, University of Bonn, Germany [Hans-Peter Fischer], Molecular Genetics of Breast Cancer, Deutsches Krebsforschungszentrum (DKFZ), Heidelberg, Germany [Ute Hamann], Institute for Prevention and Occupational Medicine of the German Social Accident Insurance, Institute of the Ruhr University Bochum (IPA), Bochum, Germany [Thomas Brüning, Beate Pesch, Sylvia Rabstein, Anne Lotz]; and Institute of Occupational Medicine and Maritime Medicine, University Medical Center Hamburg-Eppendorf, Germany [Volker Harth]. HEBCS thanks Johanna Kiiski, Carl Blomqvist, Kristiina Aittomäki, Kirsimari Aaltonen, Karl von Smitten, Irja Erkkilä. HMBCS thanks Peter Hillemanns, Hans Christiansen and Johann H. Karstens. HUBCS thanks Shamil Gantsev. ICICLE thanks Kelly Kohut, Michele Caneppele, Maria Troy. KARMA and SASBAC thank the Swedish Medical Research Counsel. KBCP thanks Eija Myöhänen. kConFab/AOCS wish to thank Heather Thorne, Eveline Niedermayr, all the kConFab research nurses and staff, the heads and staff of the Family Cancer Clinics, and the Clinical Follow Up Study (which has received funding from the NHMRC, the National Breast Cancer Foundation, Cancer Australia, and the National Institute of Health (USA)) for their contributions to this resource, and the many families who contribute to kConFab. LMBC thanks Gilian Peuteman, Thomas Van Brussel, EvyVanderheyden and Kathleen Corthouts. MABCS thanks Milena Jakimovska (RCGEB “Georgi D. Efremov”), Snezhana Smichkoska, Emilija Lazarova (University Clinic of Radiotherapy and Oncology), Mitko Karadjozov (Adzibadem-Sistina Hospital), Andrej Arsovski and Liljana Stojanovska (Re-Medika Hospital) for their contributions and commitment to this study. MARIE thanks Petra Seibold, Dieter Flesch-Janys, Judith Heinz, Nadia Obi, Alina Vrieling, Sabine Behrens, Ursula Eilber, Muhabbet Celik, Til Olchers and Stefan Nickels. MBCSG (Milan Breast Cancer Study Group): Paolo Radice, Bernard Peissel, Jacopo Azzollini, Erica Rosina, Daniela Zaffaroni, Bernardo Bonanni, Irene Feroce, Mariarosaria Calvello, Aliana Guerrieri Gonzaga, Monica Marabelli, Davide Bondavalli and the personnel of the Cogentech Cancer Genetic Test Laboratory. The MCCS was made possible by the contribution of many people, including the original investigators, the teams that recruited the participants and continue working on follow-up, and the many thousands of Melbourne residents who continue to participate in the study. We thank the coordinators, the research staff and especially the MMHS participants for their continued collaboration on research studies in breast cancer. The following are NBCS Collaborators: Anne-Lise Børresen-Dale (Prof. Em.), Kristine K. Sahlberg (PhD), Lars Ottestad (MD), Rolf Kåresen (Prof. Em.) Dr. Ellen Schlichting (MD), Marit Muri Holmen (MD), Toril Sauer (MD), Vilde Haakensen (MD), Olav Engebråten (MD), Bjørn Naume (MD), Alexander Fosså (MD), Cecile E. Kiserud (MD), Kristin V. Reinertsen (MD), Åslaug Helland (MD), Margit Riis (MD), Jürgen Geisler (MD), OSBREAC and Grethe I. Grenaker Alnæs (MSc). NBHS and SBCGS thank study participants and research staff for their contributions and commitment to the studies. For NHS and NHS2 the study protocol was approved by the institutional review boards of the Brigham and Women’s Hospital and Harvard T.H. Chan School of Public Health, and those of participating registries as required. We would like to thank the participants and staff of the NHS and NHS2 for their valuable contributions as well as the following state cancer registries for their help: AL, AZ, AR, CA, CO, CT, DE, FL, GA, ID, IL, IN, IA, KY, LA, ME, MD, MA, MI, NE, NH, NJ, NY, NC, ND, OH, OK, OR, PA, RI, SC, TN, TX, VA, WA, WY. The authors assume full responsibility for analyses and interpretation of these data. OBCS thanks Arja Jukkola-Vuorinen, Mervi Grip, Saila Kauppila, Meeri Otsukka, Leena Keskitalo and Kari Mononen for their contributions to this study. The OFBCR thanks Teresa Selander, Nayana Weerasooriya and Steve Gallinger. ORIGO thanks E. Krol-Warmerdam, and J. Blom for patient accrual, administering questionnaires, and managing clinical information. The LUMC survival data were retrieved from the Leiden hospital-based cancer registry system (ONCDOC) with the help of Dr. J. Molenaar. PBCS thanks Louise Brinton, Mark Sherman, Neonila Szeszenia-Dabrowska, Beata Peplonska, Witold Zatonski, Pei Chao, Michael Stagner. The ethical approval for the POSH study is MREC /00/6/69, UKCRN ID: 1137. We thank staff in the Experimental Cancer Medicine Centre (ECMC) supported Faculty of Medicine Tissue Bank and the Faculty of Medicine DNA Banking resource. The RBCS thanks Jannet Blom, Saskia Pelders, Wendy J.C. Prager – van der Smissen, and the Erasmus MC Family Cancer Clinic. SBCS thanks Sue Higham, Helen Cramp, Dan Connley, Ian Brock, Sabapathy Balasubramanian and Malcolm W.R. Reed. We thank the SEARCH and EPIC teams. SZBCS thanks Ewa Putresza. UCIBCS thanks Irene Masunaka. UKBGS thanks Breast Cancer Now and the Institute of Cancer Research for support and funding of the Generations Study, and the study participants, study staff, and the doctors, nurses and other health care providers and health information sources who have contributed to the study. We acknowledge NHS funding to the Royal Marsden/ICR NIHR Biomedical Research Centre.

